# Imaging breast cancer using a dual-ligand nanochain particle

**DOI:** 10.1101/411728

**Authors:** Gil Covarrubias, Anthony Cha, Abdelrahman Rahmy, Morgan Lorkowski, Vindya Perera, Bernadette O. Erokwu, Chris Flask, Pubudu M. Peiris, William P. Schiemann, Efstathios Karathanasis

**Affiliations:** Department of Biomedical Engineering, Case Western Reserve University, Cleveland, Ohio; Department of Radiology, Case Western Reserve University, Cleveland, Ohio; Case Center for Imaging Research, Case Western Reserve University, Cleveland, Ohio; Case Comprehensive Cancer Center, Case Western Reserve University, Cleveland, Ohio

**Keywords:** Iron oxide nanochains, dual-ligand nanoparticle, breast cancer, MRI

## Abstract

Nanoparticles often only exploit the upregulation of a receptor on cancer cells to enhance intratumoral deposition of therapeutic and imaging agents. However, a single targeting moiety assumes that a tumor is homogenous and static. Tumoral microenvironments are both heterogenous and dynamic, often displaying variable spatial and temporal expression of targetable receptors throughout disease progression. Here, we evaluated the *in vivo* performance of an iron oxide nanoparticle in terms of targeting and imaging of orthotropic mouse models of aggressive breast tumors. The nanoparticle, a multi-component nanochain, was comprised of 3-5 iron oxide nanoparticles chemically linked in a linear chain. The nanoparticle’s surface was decorated with two types of ligands each targeting two different upregulated biomarkers on the tumor endothelium, P-selectin and fibronectin. The nanochain exhibited improved tumor deposition not only through vascular targeting but also through its elongated structure. A single-ligand nanochain exhibited a ∼2.5-fold higher intratumoral deposition than a spherical nanoparticle variant. Furthermore, the dual-ligand nanochain exhibited higher consistency in generating detectable MR signals compared to a single-ligand nanochain. Using a 7T MRI, the dual-ligand nanochains exhibited highly detectable MR signal within 3h after injection in two different animal models of breast cancer.

## INTRODUCTION

Imaging is critical for management of patients with breast cancer including diagnosis, treatment planning and response assessments. To improve cancer imaging, various targeting schemes have been employed to direct nanoparticle imaging agents to cancers [1, 2]. Traditional targeting strategies decorate the surface of nanoparticles with a ligand directing them to upregulated receptors on breast cancer cells within the tumor interstitium. Rather than targeting the tumor interstitium, an alternative strategy is to use vascular targeting and direct the nanoparticles to the altered endothelium associated with breast cancer. The endothelium of tumors, including those of the breast, displays a wide variety of targetable biomarkers that are not readily found on healthy endothelium. For circulating nanoparticles, the endothelium is the closest point-of-contact, which facilitates direct access to the targetable vascular biomarkers of the disease [3-9]. Considering their size and multivalent avidity, nanoparticles are ideal for vascular targeting.

Further, the shape of the nanoparticle can dictate its targeting avidity [7-10-12]. Recently, we reported a new one-pot synthetic concept for making multicomponent chain-like nanoparticles (termed nanochains), which are comprised of about three iron oxide nanospheres chemically linked into a linear, chain-like assembly [13]. The chain-like shape of the nanoparticles facilitates geometrically enhanced multivalent attachment on vascular targets, resulting in rapid and effective deposition of the nanochains onto the endothelium of tumors [5, 8, 13, 14].

In addition to adjusting the shape of nanoparticles, they can also be decorated with more than one type of targeting ligand [3, 15]. This provides great flexibility in the particle’s design allowing targeted nanoparticles to consider the dynamic nature of tumors. As cancer cells evolve, the surrounding vascular reflects this behavior by having continual changes in the expression of targetable biomarkers both spatially and temporally [16-22]. In previous work [3, 6], we have shown that a multi-ligand nanoparticle with two or more types of ligands can account for the spatiotemporal alterations in the expression patterns of targetable receptors on the endothelium of tumors, which are missed by single-ligand nanoparticles.

Here, we evaluated the performance of a dual-ligand nanochain as a targeted MR imaging agent and its ability to target breast cancer. We employed two different peptides as ligands on the nanoparticles that target 1) a vascular receptor on the remodeled endothelium of tumors (P-selectin) [23-33], and 2) an extracellular biomarker in the near-perivascular regions of tumors (fibronectin) [34-37]. It should be noted that the tumor endothelium often acts as a mirror reflecting various cancerous activities of the tumor interstitium. Overexpression of P-selectin is linked with angiogenesis and is prominent on proliferating endothelial cells. On the other hand, abundance of fibronectin in the perivascular regions of tumors is strongly associated with migration and invasion of cancer cells. Considering their insignificant expression on the endothelium of healthy tissues, these biomarkers are an ideal fit to a vascular targeting scheme. Using a mouse syngeneic model of triple-negative breast cancer, we compared the vascular targeting abilities of the dual-ligand nanochain to a single-ligand nanochain variant and its spherical counterpart. Overall, we show that the combination of two different ligands on the nanochain particle effectively captures the dynamic nature of breast cancers and targets the spatial and temporal variations in receptor presentation on the tumor endothelium.

## MATERIALS AND METHODS

### Synthesis of Parent Iron Oxide Nanoparticles

Iron oxide nanoparticles are synthesized using a co-precipitation method of Fe(II) and Fe(III) ions. Briefly, a 2 to 1 molar ratio of FeCl_3_.6H_2_O and FeCl_2_.4H_2_O was dissolved in a 5mL solution of deoxygenated water. To the iron chloride solution, 5mL of a 0.4M HCl solution was added and allowed to stir vigorously. The iron precursor solution was then added to a 50mL solution of 0.5M NaOH at 80^∼^C under a constant flow of nitrogen. The reaction continued for an additional 15 minutes with stirring. Upon placing the iron precursor solution in the preheated NaOH solution, the reaction mixture immediately turned black indicating the formation of iron oxide nanoparticles. Using magnetic separation, the iron oxide solution was cleaned using deoxygenated water until a stable ferrofluid was developed. Once a stable ferrofluid was obtained, the particle volume was increased to a total volume of 50mL using deoxygenated water. In order to prevent the aggregation of iron oxide nanoparticles, 340 mg of anhydrous citric acid was added to the particle solution and the pH was consequentially increased to 5.2 using ammonia. This reaction was heated for 2 hours under a constant flow of nitrogen at 80°C. After the reaction was completed large and unreacted particles were removed using multiple centrifugation steps at 5000 rpm in 30-minute increments. Excess citric acid was removed using Amicon^®^ Ultra-15 centrifugal filters.

The surface of the iron oxide particle was modified using silane-PEG-NH_2_ or silane-PEG-COOH (2000kDa). Citric coated iron oxide nanoparticles were concentrated to 1mg/mL in Milli-Q water and the pH was adjusted to 11 using ammonia. 10 mg of silane-PEG-NH_2_ was added to citric coated iron oxides and allowed to react for 24 hours while shaking. The reaction was taken to completion by heating the nanoparticle solution to 80°C for 2 hours in order to achieve covalent linking between the iron oxide surface and the polymer. The resultant PEGylated nanoparticles (Fe_3_O_4_@silane-PEG-COOh or NH_2_), were concentrated using Amicon^®^ Ultra-15 centrifugal filters. The concentrated IONP-NH_2_ solution was stored at 4^°^C.

Fluorescent labelling was used to determine the total amount of functional groups on the surface of the iron oxide nanoparticles. To assess the quantity of amine functional groups, a 10-molar excess of Alexa Fluor^®^ 488 NHS ester (Invitrogen, Carlsbad, CA) was added to the IONP-NH_2_ particles and allowed to react for 2 hours in the dark. The excess fluorophore was dialyzed out in PBS using a 100,000 Da MW cut-off membrane bag. All fluorescent measurements were performed on a fluorescent plate reader (Synergy HT; BioTek Instruments, Winooski, VT) using a 480nm excitation and a 520 emission. The fluorescent intensity of the nanoparticles was compared to a standard curve. Iron content was measured using ICP-OES after sample digestion in a 1 to 4 volumetric ratio of concentrated HNO_3_ to concentrated HCl. Iron oxide nanoparticle calculations were performed assuming only Fe_3_O_4_ particles were made and with a 5.2 g/cm3 density.

### Synthesis of Chain-like Nanoparticles

First, the parent nanoparticles (IONP-COOH and IONP-NH_2_) were transferred from water to dimethylformamide (DMF) and heated to evaporate all water. The concentration of the nanoparticles was then adjusted to 1 mg/mL Fe. The carboxyl groups on the IONP-COOH nanoparticles were activated with an excess of *N,N’*-Dicyclohexylcarbodiimide over the available COOH groups in the presence of pyridine.

IONP-NH_2_ nanoparticles and the activated IONP-COOH were mixed at a ratio of 2.5:1 and allowed to react for 30 min. The reaction was then arrested by ‘deactivating’ the carboxyl groups using a 10-molar excess of ethylenediamine (relative to the number of carboxyl groups). Finally, nanochains were transferred to water and were separated by centrifugation with Amicon^®^ Ultra-15 centrifugal filters. To further clean nanochains from any unreacted parent nanoparticles, a strong magnet was used.

### Functionalization of Nanoparticles with Targeting Ligands

Functionalization of nanoparticles with vascular targeting ligands was accomplished via a sulfo-SMCC crosslinker. The nanoparticle composites, IONP-NH_2_ and NC-NH_2_, had terminal amine groups that are readily available for conjugation with available thiol groups on the cysteine end of the targeting ligands. Here P-selectin-targeting peptide CDAEWVDVS and fibrin-targeting peptide CREKA were used. Briefly, sulfo-SMCC contains two functional terminal groups on contralateral sides; an amine-reactive N-hydroxysuccinimide (NHS ester) and a sulfhydryl-reactive maleimide group. First, sulfo-SMCC in a 2-molar excess to available amine groups on the IONP-NH_2_ solution and allowed to react for 30 minutes while shaking. Next, a 2:3 molar excess of sulfo-SMCC to targeting ligand was added and allowed to react for an additional 2 hours while shaking. The functionalized product was dialyzed against PBS using a 100,000 Da MW cut-off dialysis bag to remove excess peptide and crosslinker. For dual-ligand functionalization equal molar peptides were added with the overall molar excess to sulfo-SMCC still at a 2:3 molar ratio (2-moles sulfo-SMCC:1.5 moles fibrin-targeting peptide:1.5 moles P-selectin targeting peptide).

Bio-Rad DC protein assay was used to quantify the total number of conjugated peptides. Here, 200 µL of Bio-Rad dye solution (1 to 2 parts Bio-Rad dye and water) was added to an 800 µL solution of 10 mg/mL particles and vortexed. The absorbance of the sample was obtained at 595 nm after incubating the sample for 15 minutes. The absorbance value was compared to a standard curve, which was obtained by measuring the known absorbance of known concentrations of CREKA, P-selectin peptide or both.

### Murine Tumor Models

All animal procedures were conducted under a protocol approved by the Institutional Animal Care and Use Committee (IACUC) at Case Western Reserve University. The well-being of the animals took priority over continuation of planned interventions. All animals received standard care, including *ad libitum* access to food and water, a 12/12 light/dark cycle, appropriate temperature and humidity. All animals received standard care ensuring proper protocol guidelines were followed.

We used two different mammary fat pad models developed from 4T1 and D2.A1 cancer cell lines. Both cell lines were transfected with both a luciferase and a green fluorescent protein (GFP) encoding lentivirus. 4T1 and D2.A1 mammary fat pad inoculations were preformed per approved institutional protocol. Briefly, 0.5 x 106 4T1-Luc-GFP cells were orthotopically inoculated in the right 9th inguinal gland of female BALB/cJ mice while anesthetized using an isoflurane inhalant. Using a previously established timeline, mice studies were preformed approximately 10-14 days post 4T1 inoculations until bioluminescent signals reached 1-2 x108 photons/sec. D2.A1 studies were performed on a similar timeline. The animals were closely monitored on a daily basis to ensure they did not suffer adverse effects resulting from tumor inoculations.

The well-being of the animals took priority over precise measurements in decisions regarding euthanasia or other interventions. All procedures were conducted using anesthetic to minimize pain and distress. The inhalant anesthetic, isoflurane, was used as the primary anesthesia in our experiments. However, developing tumors may ultimately result in some level of distress or discomfort in these mice. If, during the time following tumor inoculation the animal showed signs of post-procedure pain, the animal was euthanized. The research team, as well as the veterinary team of the animal facility, diligently monitored the condition of the animals, and removed any animal exhibiting signs of pain or distress as soon as humanly possible. When an animal showed distress or stopped eating and drinking (visually evaluated or there was a 15% loss of body weight), the animal was immediately euthanized. If it was observed that the tumor became 10% of the body mass of the animal or if there were changes in grooming, weight, behaviors, or kyphosis, the animal were immediately euthanized. Additionally, if we observed that an animal was suffering from inactivity, prostration, labored breathing, sunken eyes, hunched posture, piloerection/matted fur, unresolving skin ulcers, abnormal vocalization when handled, emaciation or anorexia, the animal was immediately euthanized. In all cases euthanasia was carried out in a CO_2_ chamber.

### Bioluminescent and *Ex Vivo* Fluorescent Imaging

Using the IVIS Spectrum system, bioluminescent imaging (BLI) was performed 10 minutes after an intraperitoneal injection of 200 µL of a 12.5 mg/mL solution of D-luciferin in sterile PBS. A BLI was taken every 3 days until the terminal point of the study. At the terminal point of the study, tumors were resected and used for *ex vivo* GFP fluorescent imaging, *ex vivo* tissue relaxivity or histological analysis.

### Histological Analysis

Immunohistochemistry was performed to determine the topological distribution of available targeting sites for the ligands of interest, fibrin-associated proteins and P-selectin. Mice were anesthetized with an intraperitoneal injection of ketamine/xylazine and transcardially perfused with heparinized PBS followed by 4% paraformaldehyde in PBS. Mammary tumor tissues were resected and placed in 4% paraformaldehyde for 24 hours. Tumor tissues were washed 3 times in PBS and placed in a 30% sucrose (w/v) for 48 hours. Finally, the tumor tissues were placed in OCT and stored at −80°C for two days prior to cryosectioning. Subsequent tissue sections were sliced at 12 µm and used for histological staining. To identify the location of fibronectin, P-selectin and the tumor microvasculature immunohistochemistry staining was performed using anti-fibrin and anti-P-selectin primary antibodies for each of the ligands of interest and an endothelial antigen CD31 for the vasculature. A secondary antibody tagged with Alexa Fluor^®^ 568 was added to do fluorescent imaging on the stained tissue sections. The tissue sections were counterstained using a DAPI nuclear stain. Tumor cells were imaged with their GFP tagged marker. The tissue sections were imaged at 5x, 10x and 20x using a Zeiss Axio Observer Z1 motorized inverted fluorescent microscope. For larger sized montages, the Axio Vision software automatic tiling was achieved using the Mosaic acquisition feature. Nanochain particles were detected using a Prussian Blue stain and imaged using brightfield.

### *Ex Vivo* Tissue Relaxivity

The *ex vivo* tissue relaxivity was calculated using a 1.5 T Bruker Minispec mq60. Mice were anesthetized with an intraperitoneal injection of ketamine/xylazine and transcardially perfused with heparinized PBS. Kidneys, liver, spleen, lungs and tumors were excised and grinded in 1 mL of water. Grounded tissues were placed in 5mm disposable grade NMR tubes. Samples were placed in the 1.5 T Bruker Minispec and relaxation values were recorded for use in the analysis of tissue relaxivity per mass of collected tissue.

### MR imaging

MR images were acquired using a 7 T Bruker MRI system. A volume coil (3.5 cm inner diameter) was employed. The sequence used was a Rapid Acquisition with Relaxation Enhancement (RARE). High-resolution images were obtained before and 3 hours after IV injection of the nanochains (at a dose of 10 mg Fe per kg b.w.) using a *T*2-weighted RARE sequence with the following parameters: TR/TE = 3646.6/31 ms, matrix = 256 × 256, FOV = 3 × 3 cm, and 5 averages. The acquisition time approximately 10 minutes using a gating acquisition method. This resulted in an in-plane spatial resolution of 111.7 μm and a slice thickness of 1 mm.

### Statistical Analysis

Statistics were performed in Prism version 7 for Mac (GraphPad Software, La Jolla, CA, USA). All the experiments were performed in triplicates unless stated otherwise. Data are represented as mean±s.d. In cases where data met the assumptions necessary for parametric statistics, analysis of differences between two groups was performed using two-tailed Student’s t-test assuming equal variance. Data from three or more groups were analyzed with a one-way analysis of variance (ANOVA) that was corrected for multiple comparisons using the Holm-Sidak method.

To determine the detection accuracy of particle conjugates z-score probabilities values were obtained to evaluate the effective belonginess of a single particle conjugate member to a healthy mammary fat pad population. The z-score probabilities were obtained via a confidence interval of 90% (α was set to 0.1). If a population member fell inside of the 90% confidence interval it was considered to be part of the healthy mammary fat pad population; these members were considered to be false negatives as the tumors were present however the p-values indicated they fell in the healthy population.

## RESULTS

### Nanoparticle Fabrication

We recently developed a new simple ‘one-pot’ synthetic concept for making iron oxide nanochains with high yield and consistency [13]. Fig 1a shows an illustration of the nanochain particles. Briefly, the one-pot synthesis utilized two types of the parent iron oxide nanoparticle (IONP) based on the functional group on the particle’s surface (Fig 1b). The IONP were decorated with either PEG-amine (NP-NH_2_) or PEG-carboxyl (NP-COOH). First, the carboxyl groups on the NP-COOH particles were activated with DCC. To avoid hydrolysis of the activated COOH intermediate, the one-pot synthesis was performed in an organic solvent (*i.e.*, absence of water). Once activated NP-COOH and NP-NH_2_ were mixed, the nanoparticles started reacting with each other. The reaction rate and size of the agglomerates can be dictated by the stirring rate, stoichiometry and concentration of the starting IONP particles. By mixing NP-NH_2_ and NP-COOH at a ratio of ∼2:1, two NP-NH_2_ particles were initially linked with one ‘activated’ NP-COOH forming a trimeric nanochain. We identified the optimal reaction time and parameters that generated well-defined short nanochains and not large agglomerates. Using dynamic light scattering, longitudinal measurement of the hydrodynamic size of the reaction mixture indicated the progression and growth of the nanochain. Fig 1c shows that the two populations of the starting parent IONP with sizes of 21 and 33 nm disappeared, while the nanochains appeared in a new population with the mean size being ∼80 nm. The size and structure of the nanochains was confirmed in TEM images (Fig 1d). The iron concentration was measured using ICP-OES, which was used in all the *in vivo* studies to accurately calculate the dose of the agents that was injected into the animals. Based on previous work [13], we also quantified the number of primary amines on the surface of nanochains using fluorescence labelling. To determine the number of surface amines, an excess of Alexa Fluor^®^ 488 NHS ester reacted with the nanoparticles for 2h followed by extensive dialysis to remove unbound fluorophore. Fluorescence measurements showed that the nanochain exhibited about 700 amines per particle. The fibronectin-targeting peptide (CREKA) and P-selectin-targeting peptide CDAEWVDVS were effectively conjugated onto the available amines on the surface of the nanochains using the heterobifunctional crosslinker sulfo-SMCC. Using HPLC assays, the number of peptides was quantified confirming that the single-ligand nanochains displayed ∼700 peptides per particle, whereas the dual-ligand variant had the available amines split approximately in half for each peptide. More details on the synthesis and characterization of the nanochains can be found in a previous publication [13].

**Fig 1.**
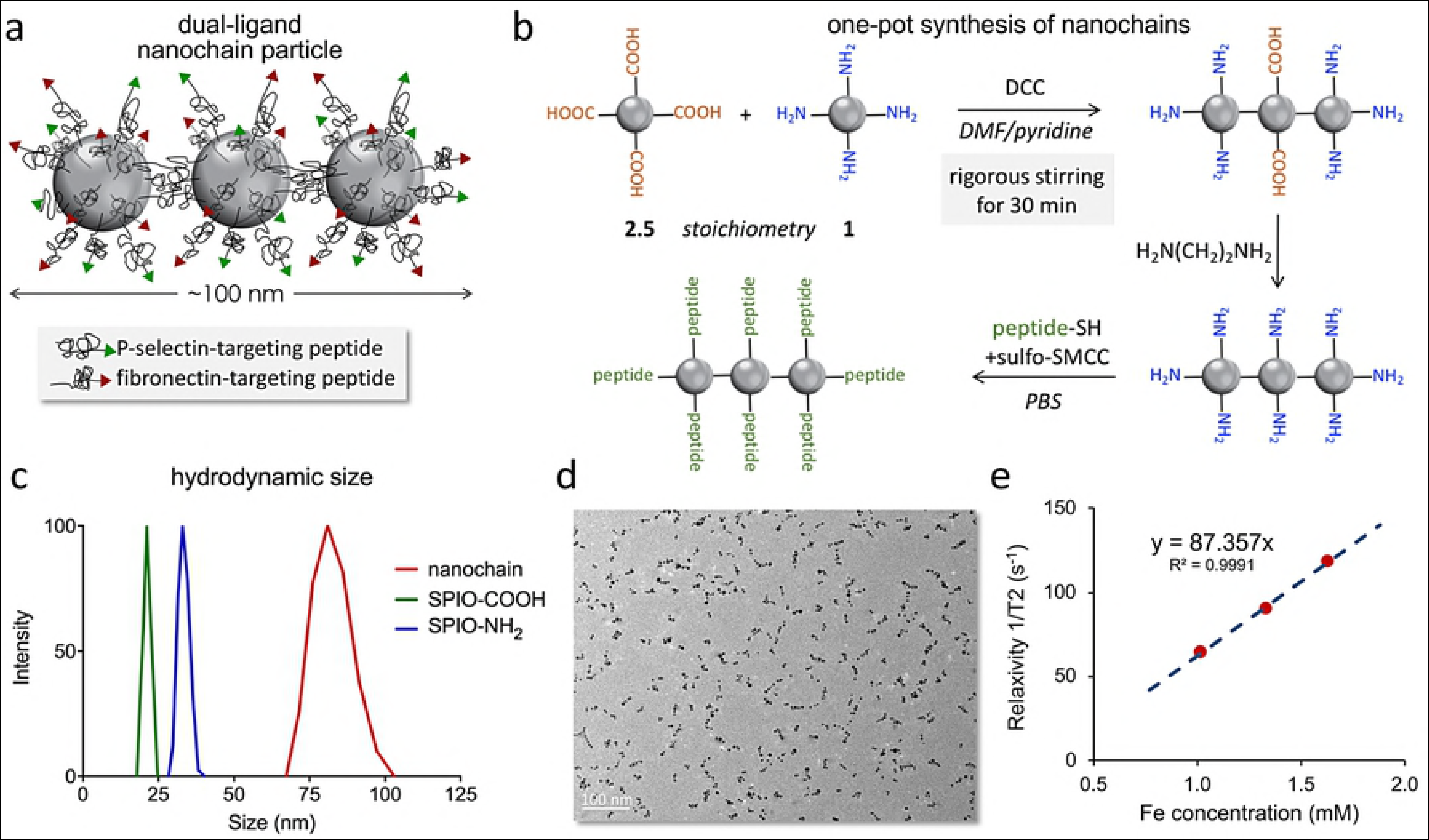
Synthesis and characterization of the nanochain particles. (a) Illustration shows the dual-ligand nanochain particle. (b) Reaction scheme shows the synthesis of nanochains using parent iron oxide nanoparticles with different surface functionality. (c) The size of the parent nanoparticles and nanochains was measured using dynamic light scattering (DLS). (d) TEM image of nanochain particles is shown. (e) The transverse (R2) relaxivity of the nanochains was measured at 1.5 Tesla using a relaxometer.

### Organ distribution

We evaluated the biodistribution of the dual-ligand nanochains in the liver, spleen, lungs and kidneys of mice 3 h after systemic administration *via* tail vein injection. The 3-hour time point was selected based on the timeframe of vascular targeting of nanoparticles as explained in the next section. The organ distribution of the dual-ligand nanochain was compared to a single-ligand nanochain, the non-targeted nanochain variant, and the parent IONP with a single ligand. In previous studies [13], the tissue deposition of targeted iron oxide nanoparticles was measured using ICP-OES, which provided direct measurement of iron concentration in tissues using ICP. However, the MR signal from tissues does not always correspond to the exact concentration of the iron oxide particles in tissues. Iron oxide particles exhibit complex relaxation properties *in vivo*, which is significantly influenced by iron clustering in the case of intracellular internalization of the nanoparticles. Since the intended use of the dual-ligand nanochain is as an MR imaging agent, we elected to measure the R2 relaxation rate in tissue as a surrogate metric of MR signal with higher R2 value indicating stronger signal. Briefly, 3 h after injection of the nanoparticles, mice were transcardially perfused with PBS and each organ was excised and homogenized. The relaxation of samples of the homogenate was measured using a 1.5 T Bruker Minispec and was recorded as tissue relaxivity per gram of tissue. This provided a robust method to measure multiple samples in a convenient and quantitative manner. Fig 2 shows a comparison of the organ distribution of the different nanoparticle variants. We tested dual-ligand nanochains targeting fibronectin and P-selectin, non-targeted nanochains, single-ligand nanochains targeting P-selectin, and single-ligand parent IONP targeting P-selectin. All the nanoparticles displayed similar biodistribution patterns with the majority of the particles being cleared by the reticuloendothelial organs (liver and spleen). The only difference was the higher clearance of the spherical IONP from the spleen compared to all the nanochain formulations. This was in good agreement with previous observations [5, 13, 38]. Further, the signal from the lungs and kidneys was negligible for all formulations. These findings indicate that the use of one or two ligands did not alter significantly the overall biodistribution patterns of the nanochains.

**Fig 2.**
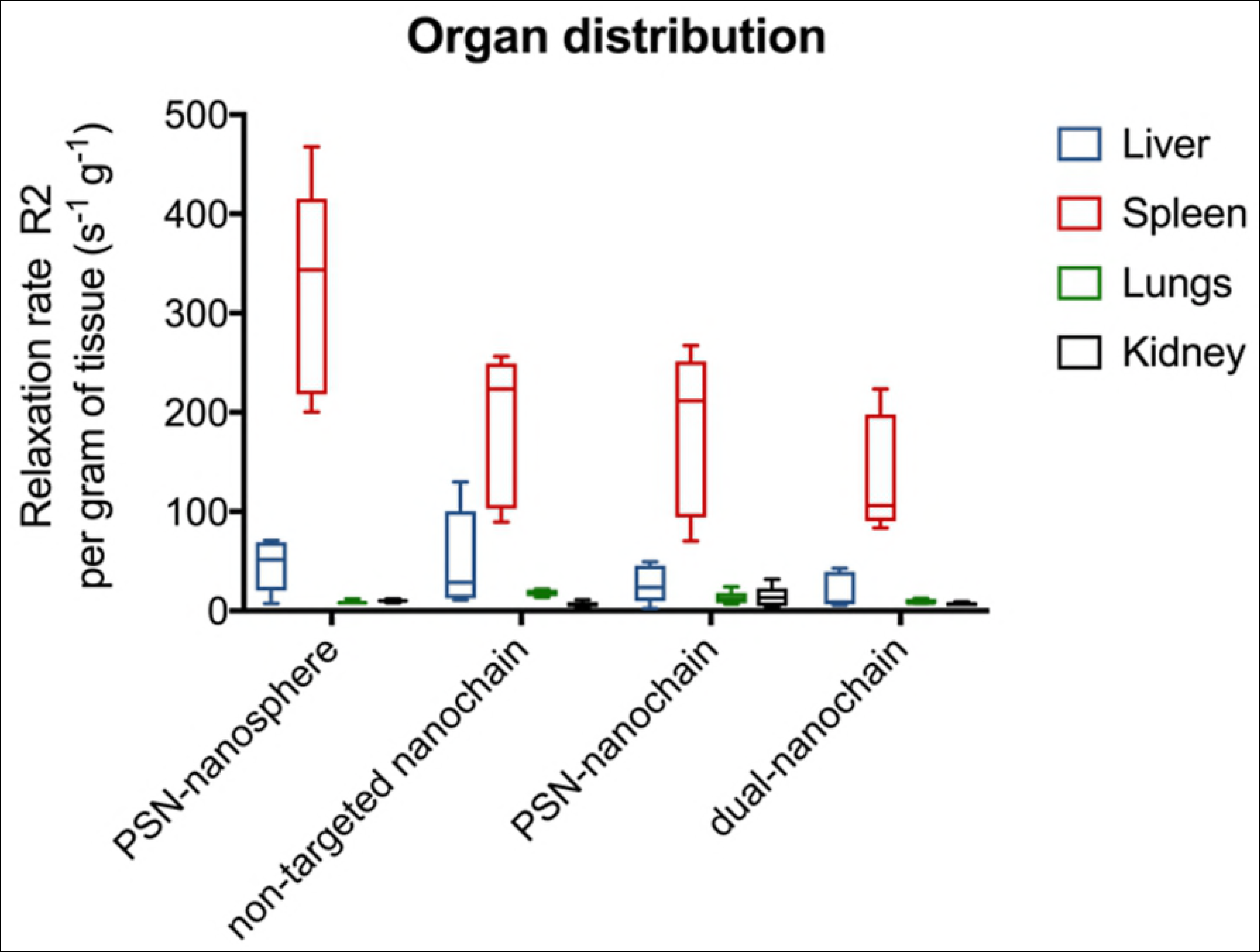
Organ distribution of targeting variants of spherical or chain-like iron oxide nanoparticles. Mice were systemically injected with nanoparticles at a dose of 10 mg/kg Fe. The dual-ligand nanochain was compared to non-targeted nanochain, single-ligand nanochain targeting P-selectin and single-ligand spherical nanoparticles targeting P-selectin (n=5 mice per formulation). Animals were euthanized 3 h after injection and organs were extracted and homogenized. The relaxation of the homogenate of different organs was measured using a 1.5 T Bruker Minispec and was recorded as tissue relaxivity per gram of tissue.

### Tumor targeting

For these studies, we used the D2.A1 cancer cell line, which is a syngeneic mouse model of triple-negative breast cancer. The D2.A1 cell line was engineered to stably express firefly luciferase and green fluorescent protein (GFP). Using this cell line, we developed an orthotopic D2.A1 model by inoculating the cancer cells into the mammary fat pad of BALB/c mice. Being a very aggressive breast cancer, the typical survival of the animals is about 25 days. Bioluminescent imaging was conducted ever 2-3 days post inoculation to monitor tumor growth. Animals were selected for the targeting studies when bioluminescence signal of the tumor reached a value of about 1.5 x 108 photons/s (approximately 2 weeks after tumor inoculation).

First, we compared the targeting performance of a single-ligand nanochain to its spherical IONP counterpart. It should be noted that the spherical nanoparticles were the parent particles used for the fabrication of the nanochains. Our previous studies showed that deposition of targeted nanoparticles onto the endothelium of tumors is rapid and is maximized within 3 h after systemic administration [3, 5, 6, 13]. Thus, we selected the 3-hour time point as the terminal point when the tumors were perfused, excised, and homogenized. The relaxation of the tumor homogenate was measured using a 1.5 T Bruker Minispec. As expected, when compared to nanospheres, the nanochains favored tumor deposition due to geometrically enhanced targeting of the tumor endothelium. Fig 3a shows that the single-ligand nanochains targeting P-selectin exhibited about a 2.5-fold higher intratumoral deposition than their spherical counterpart. This is in good agreement with a previous study that showed that nanochains facilitated superior vascular targeting of brain tumors than spherical nanoparticles [13]. We then compared a single-ligand nanochain to the dual-ligand nanochain. In a previous study, we showed that P-selecting-targeting nanoparticles had similar intratumoral deposition to fibronectin-targeting nanoparticles [6]. Thus, we tested only one of the two possible single-ligand nanochain variants. As shown in Fig 3b, the dual-ligand nanochains did not significantly outperform their single-ligand variants. However, comparisons of the average values of each group often does not reveal the entire diagnostic performance. For example, the single-ligand formulation could not consistently discriminate cancerous tissues even though we carefully selected animals with similar tumor burdens (Fig 3c). On the other hand, the dual-ligand formulation exhibited 100% success in separating tumor from healthy mammary tissues (Fig 3d). In previous studies, we have observed a similar consistency of targeting accuracy of multi-ligand nanoparticles in brain tumors [13] and metastasis [3].

**Fig 3.**
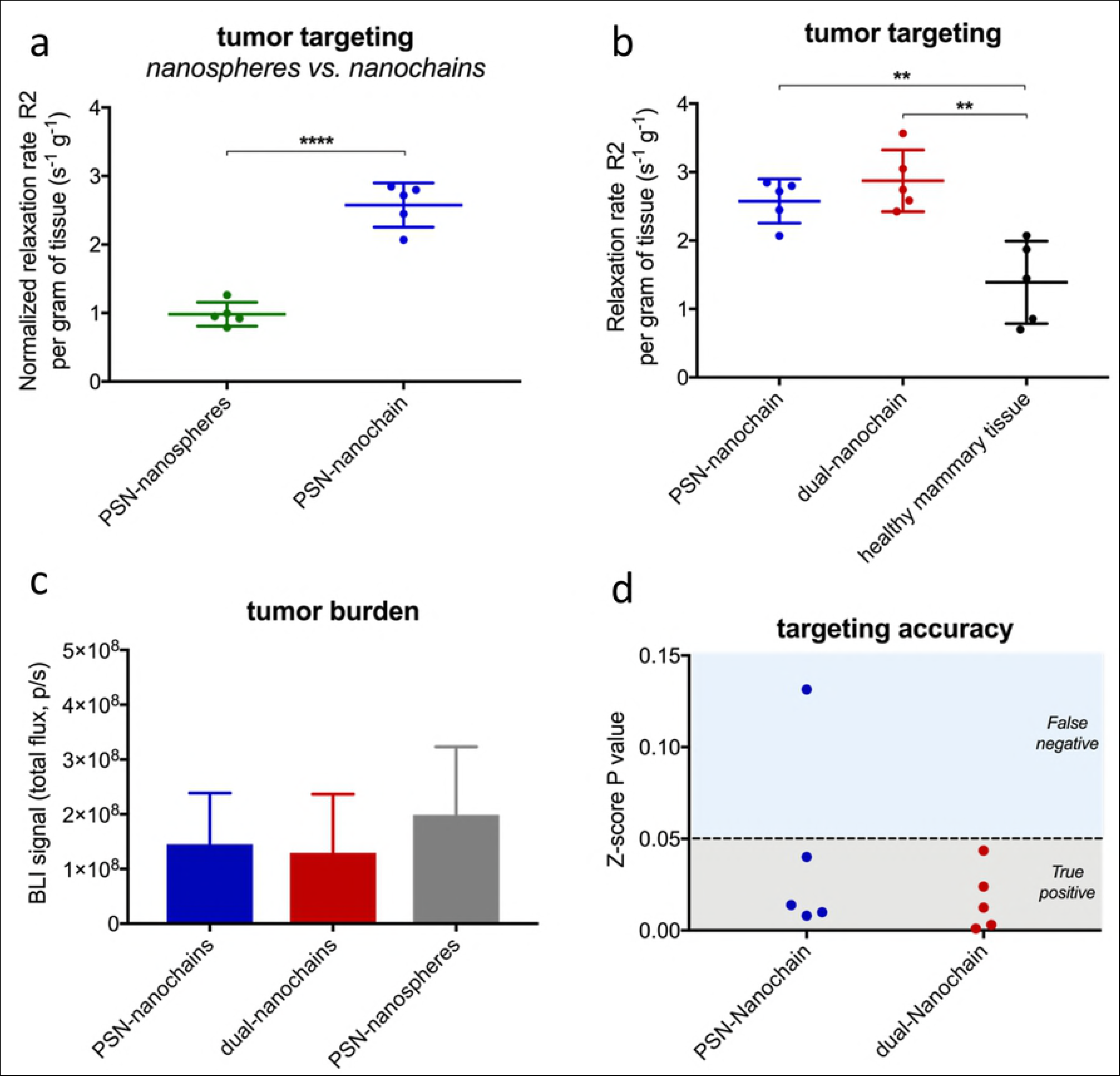
Comparison of tumor targeting accuracy of different targeted nanoparticles. (a) Single-ligand nanochains targeting P-selectin and their spherical counterparts were IV injected in mice bearing mammary D2.A1 tumors. All formulations were administered at the same dose (10 mg/kg Fe). The animals were perfused 3 h after injection and the tumors were excised, homogenized and measured using a 1.5 T Bruker Minispec (data are represented as mean?±?s.d.; n=5 mice in each group; unpaired t-test, P values: ****<0.0001). (b) The endogenous relaxation rate (R2) of healthy mammary tissue were compared to those of tumor bearing mice after systemic injection of single-ligand or dual-ligand nanochains (data are represented as mean?±?s.d.; n=5 mice in each group; unpaired t-test, P values: **<0.005). (c) Quantification of the bioluminescent signal indicated similar tumor burden of the different groups of animals. (d) Z-score analysis was performed to identify the belongingness of each tumor bearing mouse injected with a formulation to a healthy mammary population. If the z-score probability value was larger than an alpha of 0.05, it signified that it was likely that the R2 value belonged to that of a healthy mammary tissue as opposed to a diseased mammary.

### Imaging of breast tumors using MRI

The ability of the dual-ligand nanochain to image breast cancer was tested using a 7T MRI. In addition to the orthotopic D2.A1 mouse model, we also used the orthotopic 4T1 mouse model (n=3 for each tumor model). The 4T1 model is one of the standard models to study TNBC development in immunocompetent mice. MR imaging was performed before and 3 h after tail-vein injection of the dual-ligand nanochains at a dose of 10 mg/kg Fe. We used the same scanning parameters in the pre-and post-injection images. The pre-and 3h post-injection images were compared quantitatively by measuring the absolute MR signal intensity in the tumor and healthy mammary fat pad. ?he signal intensity was normalized to the signal of the healthy mammary tissue (scale: 0-1). A low value indicates high contrast in T2 images, whereas a value of 1 indicates no contrast compared to the pre-injection image. Fig 4a summarizes the measurements. The pre-injection values of the tumors in the D2.A1 and 4T1 mouse models were close to 1 indicating negligible contrast compared to health mammary. On the other hand, injection of the dual-ligand nanochains generated significant contrast enhancement in the tumors with a normalized value of 0.82 and 0.59 for the 4T1 and D2.A1 tumors, respectively. Fig 4b shows representative sagittal T2-weighted images before and 3 h after injection of the dual-ligand nanochains in the D2.A1 mouse model.

**Fig 4.**
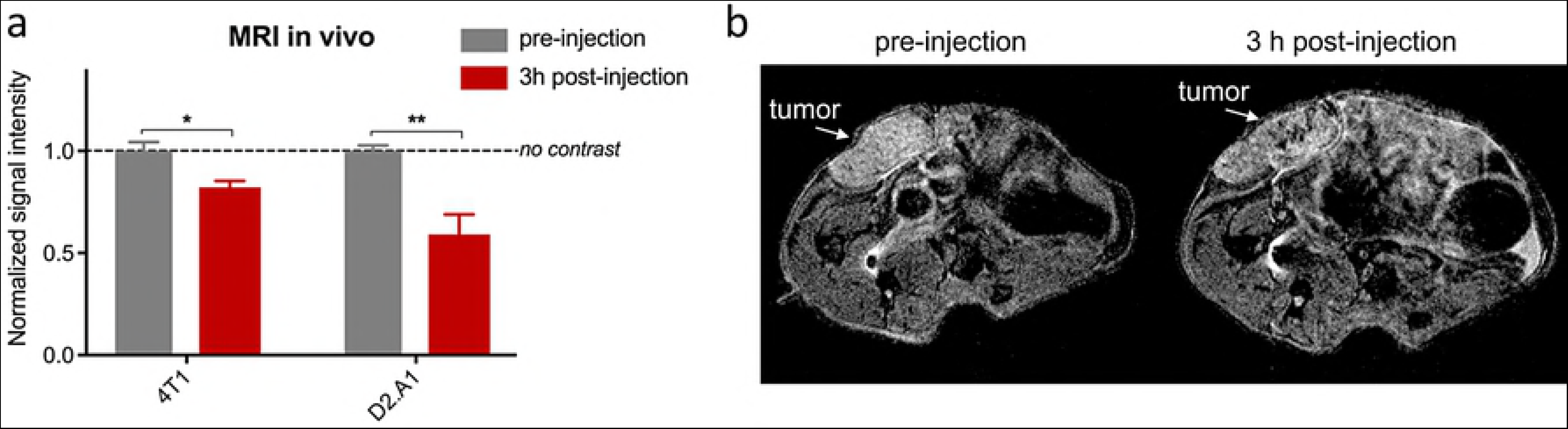
MR imaging of mice bearing mammary breast tumors using dual-ligand nanochains and a 7T MRI. (a) Animals with 4T1 or D2.A1 mammary tumors were imaged before and 3 h after tail vein injection of dual-ligand nanochains at a dose of 10 mg/kg Fe. The signal intensity was measured in the tumor and healthy mammary fat pad. ?he signal intensity normalized to the signal of the healthy mammary tissue (scale: 0-1). The normalized values of 0 and 1 correspond to maximum and minimum contrast, respectively, compared to the pre-injection values of healthy mammary tissue (data are represented as mean?±?s.d.; n=3 mice for each animal model; unpaired t-test, P values: *<0.05, **<0.005). (b) Representative sagittal T2-weighted images of a mouse bearing a D2.A1 mammary tumor before and 3 h after injection of dual-ligand nanochains.

### Histological evaluation

We performed histological analysis to confirm vascular targeting and deposition of the dual-ligand nanochains in tumors. After the last MRI session, mice were intracardially perfused with heparinized PBS. Tumors and normal mammary tissues were collected and processed for histological evaluation. Direct fluorescence imaging of GFP (green) indicated the location of breast cancer cells. The tissue slices were also immunohistochemically stained for the endothelial antigen CD31, anti-fibronectin antibody and anti-P-selectin antibody. Using fluoresence microscopy, Fig 5a shows representative images of the tumor indicating that the endothelium and near-vascular regions exhibited overexpression of fibronectin and P-selectin. We should note that negligible expression of fibronectin or P-selectin was observed on the endothelium in healthy mammary tissues (images not shown). The associated expression of P-selectin and endothelium with respect to tumor endothelium should favor vascular targeting of the dual-ligand nanochains. In addition to fluorescence microscopy, bright-field microscopy was performed on the same histological sections using the Prussian blue stain to visualize the iron oxide nanochains. Fig 5b shows that dual-ligand nanochains were predominantly distributed around blood vessels in the tumor.

**Fig 5.**
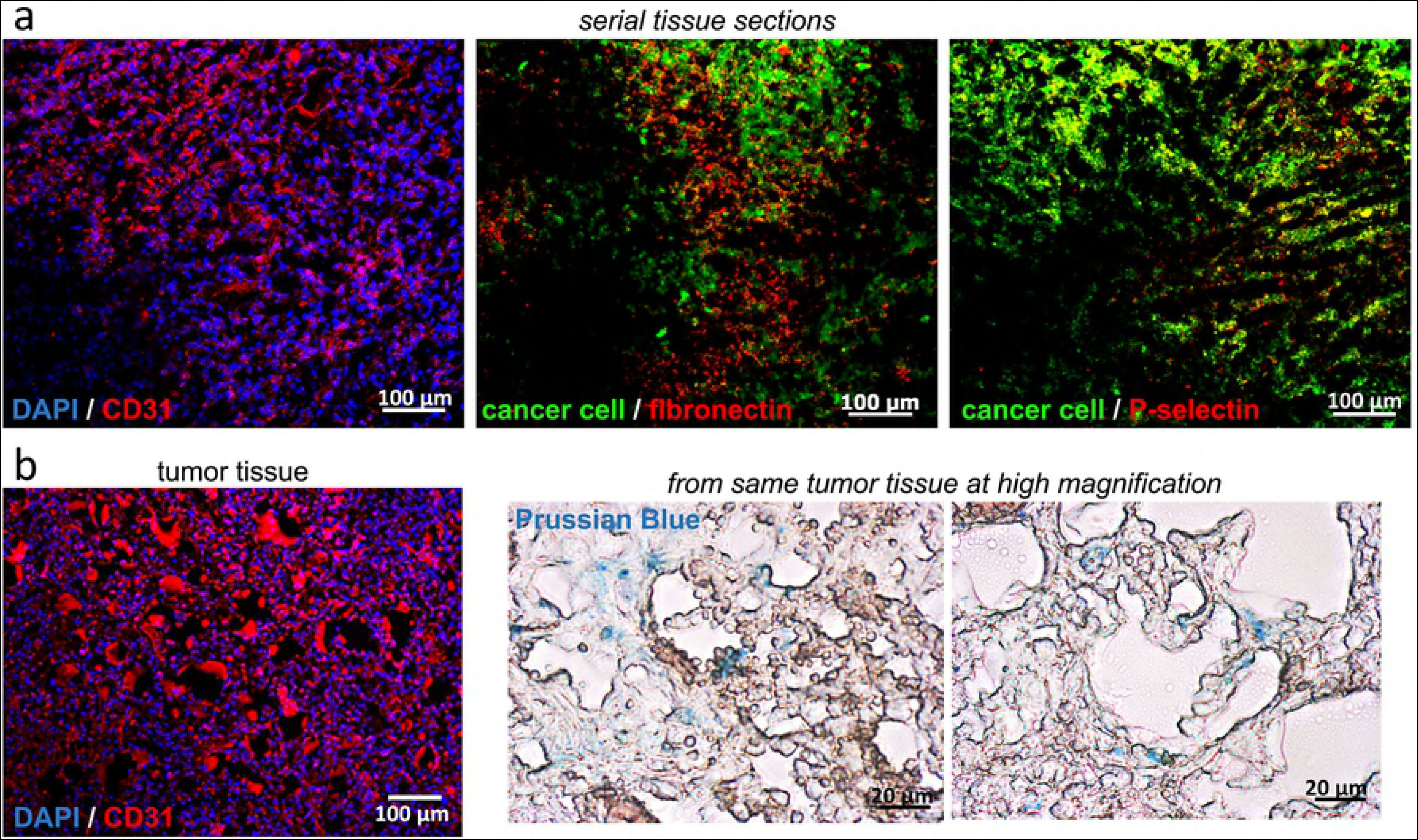
Histological evaluation of the expression of vascular biomarkers and intratumoral deposition of targeted nanoparticles in the orthotopic D2.A1 mouse model. (a) Mice bearing mammary D2.A1 tumors were euthanized 3 h after tail vein injection of dual-ligand nanochains at a dose of 10 mg/kg iron. After perfusion, tumors were excised and processed for histology. Images from serial tissue sections show that topology of fibronectin and P-selectin with respect to the endothelium (10x magnification; nuclear stain: blue; cancer cells: green; CD31 endothelial marker or fibronectin or P-selectin: red; overlay: yellow). (b) The near-perivascular deposition of the dual-ligand nanochain was identified through an iron stain (left panel: 10x magnification; nuclear stain: blue and CD31 endothelial marker: red; middle and right panels: 20x magnification; iron stain: Prussian blue).

## DISCUSSION

Due to their size and multivalent avidity, nanoparticles are ideal for vascular targeting of upregulated biomarkers on the tumor endothelium. Since the endothelium is the closest point-of-contact, circulating nanoparticles in the bloodstream have direct access to scavenge the endothelium for vascular biomarkers of cancer. Recently, we reported a new one-pot synthetic concept for making multicomponent chain-like nanoparticles (termed nanochains), which are comprised of about three iron oxide nanospheres chemically linked into a linear assembly [13]. Here we showed that targeting avidity can be dictated by adjusting the shape. Within 3 h post-injection, vascular targeting of nanochains targeting P-selectin resulted in a 2.5-fold higher deposition in breast tumors than targeting equivalent spherical nanoparticles.

Traditional targeting strategies use a single-ligand system that considers cancer as a monolithic disease and fails to account for tumor heterogeneity. However, aggressive breast tumors exhibit a dynamic tumor microenvironment with targetable vascular biomarkers being continuously changing over time and space [39]. In this context, our previous studies showed that vascular biomarkers often exhibited spatiotemporal variability, representing different stages of tumor development [3]. Considering then that the altered endothelium associated with breast cancer displays a diverse and dynamic set of targetable biomarkers. a combination of different ligands on the same particle may be required to efficiently direct a nanoparticle to the majority of a tumor volume.

In previous studies [3-9], we tested different peptides to direct nanoparticles to different vascular biomarkers that represent various microenvironments of breast cancer, including α_v_β_3_ integrin, P-selectin, EGFR, PTPµ, and fibronectin. In this work, we selected to target P-selectin and fibronectin that represent different cancerous activities and stages of breast cancer development. Upregulation of P-selectin is pronounced on proliferating endothelial cells and is associated with the early response of cancer cells to hypoxia development and angiogenesis. On the other hand, perivascular overexpression of fibronectin is critical in the migration and invasion of cancer cells [34-37]. Not surprising, the dual-ligand nanochain variants exhibited significant intratumoral deposition. We selected iron oxide as our nanomaterial basis due to its ability to generate significant contrast in MR imaging. In two different animal models of breast cancer, MR imaging and the dual-ligand nanochain facilitated precise detection of breast cancer. Taking under consideration that diagnosis, treatment planning and response assessments of breast cancer rely heavily on imaging, a more accurate imaging test for breast cancer can change patient management by guiding early therapeutic interventions before the disease becomes unmanageable.

## CONCLUSIONS

The multicomponent nanochain represents a higher order nanostructure that is comprised of individual nanoparticles. By utilizing a simple ‘one-pot’ synthesis, nanochains were fabricated with high reproducibility, yield and consistency across batches. Further, the combination of two different ligands on the same nanochain particle effectively targeted the dynamic nature of breast tumors and generated highly detectable MR signals.

## ACKNOWLEDGEMENTS

This work was partially supported by grants from the National Cancer Institute (R01CA177716, U01CA198892), the Alex’s Lemonade Stand Foundation and the Angie Fowler AYA Cancer Research Initiative of the Case Comprehensive Cancer Center (E.K.). G.C. was supported by a fellowship from the NIH Interdisciplinary Biomedical Imaging Training Program (T32EB007509) administered by the Department of Biomedical Engineering, Case Western Reserve University.

## References

1. Zhou Z, Lu ZR. Molecular imaging of the tumor microenvironment. Adv Drug Deliv Rev. 2017;113:24-48. Epub 2016/08/09. doi:10.1016/j.addr.2016.07.012. PubMed PMID: 27497513.

2. Toy R, Bauer L, Hoimes C, Ghaghada KB, Karathanasis E. Targeted nanotechnology for cancer imaging. Adv Drug Deliv Rev. 2014;76:79–97. doi:10.1016/j.addr.2014.08.002. PubMed PMID: 25116445; PubMed Central PMCID: PMC4169743.

3. Doolittle E, Peiris PM, Doron G, Goldberg A, Tucci S, Rao S, et al. Spatiotemporal Targeting of a Dual-Ligand Nanoparticle to Cancer Metastasis. ACS Nano. 2015;9(8):8012–21. doi:10.1021/acsnano.5b01552. PubMed PMID: 26203676; PubMed Central PMCID: PMC4579532.

4. Toy R, Hayden E, Camann A, Berman Z, Vicente P, Tran E, et al. Multimodal in vivo imaging exposes the voyage of nanoparticles in tumor microcirculation. ACS Nano. 2013;7(4):3118–29. doi:10.1021/nn3053439. PubMed PMID: 23464827; PubMed Central PMCID: PMC3640526..

5. Peiris PM, Toy R, Doolittle E, Pansky J, Abramowski A, Tam M, et al. Imaging metastasis using an integrin-targeting chain-shaped nanoparticle. ACS Nano. 2012;6(10):8783–95. doi:10.1021/nn303833p. PubMed PMID: 23005348.

6. Peiris PM, He F, Covarrubias G, Raghunathan S, Turan O, Lorkowski M, et al. Precise targeting of cancer metastasis using multi-ligand nanoparticles incorporating four different ligands. Nanoscale. 2018;10(15):6861-71. Epub 2018/04/06. doi:10.1039/c8nr02513d. PubMed PMID: 29620124; PubMed Central PMCID: PMCPMC5908762.

7. Toy R, Peiris PM, Ghaghada KB, Karathanasis E. Shaping cancer nanomedicine: the effect of particle shape on the in vivo journey of nanoparticles. Nanomedicine (Lond). 2014;9(1):121–34. doi:10.2217/nnm.13.191. PubMed PMID: 24354814.

8. Peiris PM, Toy R, Abramowski A, Vicente P, Tucci S, Bauer L, et al. Treatment of cancer micrometastasis using a multicomponent chain-like nanoparticle. J Control Release. 2014;173:51–8. doi:10.1016/j.jconrel.2013.10.031. PubMed PMID: 24188960.

9. Peiris PM, Deb P, Doolittle E, Doron G, Goldberg A, Govender P, et al. Vascular Targeting of a Gold Nanoparticle to Breast Cancer Metastasis. J Pharm Sci. 2015;104(8):2600–10. doi:10.1002/jps.24518. PubMed PMID: 26036431.

10. Gavze E, Shapiro M. Motion of inertial spheroidal particles in a shear flow near a solid wall with special application to aerosol transport in microgravity. Journal of Fluid Mechanics. 1998;371:59±79.

11. Lee SY, Ferrari M, Decuzzi P. Shaping nano-/micro-particles for enhanced vascular interaction in laminar flows. Nanotechnology. 2009;20 495101 (11pp).

12. Gentile F, Chiappini C, Fine D, Bhavane RC, Peluccio MS, Cheng MM, et al. The effect of shape on the margination dynamics of non-neutrally buoyant particles in two-dimensional shear flows. J Biomech. 2008;41(10):2312-8. Epub 2008/06/24. doi:S0021-9290(08)00150-4 [pii] 10.1016/j.jbiomech.2008.03.021. PubMed PMID: 18571181.

13. Perera VS, Covarrubias G, Lorkowski M, Atukorale P, Rao A, Raghunathan S, et al. One-pot synthesis of nanochain particles for targeting brain tumors. Nanoscale. 2017;9(27):9659–67. doi:10.1039/c7nr02370g. PubMed PMID: 28675230; PubMed Central PMCID: PMC5557407.

14. Peiris PM, Abramowski A, Mcginnity J, Doolittle E, Toy R, Gopalakrishnan R, et al. Treatment of invasive brain tumors using a chain-like nanoparticle. Cancer Research. 2015;75(7):1356–65.

15. Saul JM, Annapragada AV, Bellamkonda RV. A dual-ligand approach for enhancing targeting selectivity of therapeutic nanocarriers. J Control Release. 2006;114(3):277-87. Epub 2006/08/15. doi:S0168-3659(06)00250-1 [pii] 10.1016/j.jconrel.2006.05.028. PubMed PMID: 16904220.

16. Gay LJ, Felding-Habermann B. Contribution of platelets to tumour metastasis. Nat Rev Cancer. 2011;11(2):123-34. Epub 2011/01/25. doi:nrc3004 [pii] 10.1038/nrc3004. PubMed PMID:21258396.

17. Fukumura D, Jain RK. Tumor microenvironment abnormalities: causes, consequences, and strategies to normalize. Journal of cellular biochemistry. 2007;101(4):937-49. PubMed PMID:17171643.

18. Hobbs SK, Monsky WL, Yuan F, Roberts WG, Griffith L, Torchilin VP, et al. Regulation of transport pathways in tumor vessels: role of tumor type and microenvironment. Proc Natl Acad Sci U S A. 1998;95(8):4607-12. PubMed PMID:9539785; PubMed Central PMCID:PMC22537.

19. Karathanasis E, Suryanarayanan S, Balusu SR, McNeeley K, Sechopoulos I, Karellas A, et al. Imaging nanoprobe for prediction of outcome of nanoparticle chemotherapy by using mammography. Radiology. 2009;250(2):398-406. PubMed PMID:19188313.

20. Huebschman ML, Lane NL, Liu H, Sarode VR, Devlin JL, Frenkel EP. Molecular heterogeneity in adjacent cells in triple-negative breast cancer. Breast cancer. 2015;7:231–7. doi:10.2147/BCTT.S87041. PubMed PMID:26316815; PubMed Central PMCID:PMC4540115.

21. Norton KA, Popel AS, Pandey NB. Heterogeneity of chemokine cell-surface receptor expression in triple-negative breast cancer. American journal of cancer research. 2015;5(4):1295-307. PubMed PMID:26101698; PubMed Central PMCID:PMC4473311.

22. Martinez-Revollar G, Garay E, Martin-Tapia D, Nava P, Huerta M, Lopez-Bayghen E, et al. Heterogeneity between triple negative breast cancer cells due to differential activation of Wnt and PI3K/AKT pathways. Exp Cell Res. 2015;339(1):67–80. doi:10.1016/j.yexcr.2015.10.006. PubMed PMID:26453937.

23. Schnell O, Krebs B, Carlsen J, Miederer I, Goetz C, Goldbrunner RH, et al. Imaging of integrin alpha(v)beta(3) expression in patients with malignant glioma by [18F] Galacto-RGD positron emission tomography. Neuro-oncology. 2009;11(6):861–70. doi:10.1215/15228517-2009-024. PubMed PMID:19401596; PubMed Central PMCID:PMC2802406.

24. Reardon DA, Nabors LB, Stupp R, Mikkelsen T. Cilengitide: an integrin-targeting arginine-glycine-aspartic acid peptide with promising activity for glioblastoma multiforme. Expert opinion on investigational drugs. 2008;17(8):1225–35. doi:10.1517/13543784.17.8.1225. PubMed PMID:18616418; PubMed Central PMCID:PMC2832832.

25. Danhier F, Vroman B, Lecouturier N, Crokart N, Pourcelle V, Freichels H, et al. Targeting of tumor endothelium by RGD-grafted PLGA-nanoparticles loaded with paclitaxel. J Control Release. 2009;140(2):166-73. Epub 2009/08/25. doi:S0168-3659(09)00547-1 [pii] 10.1016/j.jconrel.2009.08.011. PubMed PMID:19699245.

26. Reddy GR, Bhojani MS, McConville P, Moody J, Moffat BA, Hall DE, et al. Vascular targeted nanoparticles for imaging and treatment of brain tumors. Clin Cancer Res. 2006;12(22):6677–86. doi:10.1158/1078-0432.CCR-06-0946. PubMed PMID:17121886.

27. Shamay Y, Elkabets M, Li H, Shah J, Brook S, Wang F, et al. P-selectin is a nanotherapeutic delivery target in the tumor microenvironment. Science translational medicine. 2016;8(345):345ra87. doi:10.1126/scitranslmed.aaf7374. PubMed PMID:27358497; PubMed Central PMCID:PMC5064151.

28. Laubli H, Borsig L. Selectins promote tumor metastasis. Seminars in cancer biology. 2010;20(3):169–77. doi:10.1016/j.semcancer.2010.04.005. PubMed PMID:20452433.

29. Kim YJ, Borsig L, Han HL, Varki NM, Varki A. Distinct selectin ligands on colon carcinoma mucins can mediate pathological interactions among platelets, leukocytes, and endothelium. Am J Pathol. 1999;155(2):461–72. doi:10.1016/S0002-9440(10)65142-5. PubMed PMID:10433939; PubMed Central PMCID:PMC1866847.

30. Ludwig RJ, Boehme B, Podda M, Henschler R, Jager E, Tandi C, et al. Endothelial P-selectin as a target of heparin action in experimental melanoma lung metastasis. Cancer Res. 2004;64(8):2743-50. PubMed PMID:15087389.

31. Nierodzik ML, Karpatkin S. Thrombin induces tumor growth, metastasis, and angiogenesis: Evidence for a thrombin-regulated dormant tumor phenotype. Cancer Cell. 2006;10(5):355–62. doi:10.1016/j.ccr.2006.10.002. PubMed PMID:17097558.

32. Borsig L, Wong R, Feramisco J, Nadeau DR, Varki NM, Varki A. Heparin and cancer revisited: mechanistic connections involving platelets, P-selectin, carcinoma mucins, and tumor metastasis. Proc Natl Acad Sci U S A. 2001;98(6):3352–7. doi:10.1073/pnas.061615598. PubMed PMID:11248082; PubMed Central PMCID:PMC30657.

33. Mousa SA, Petersen LJ. Anti-cancer properties of low-molecular-weight heparin: preclinical evidence. Thromb Haemost. 2009;102(2):258–67. doi:10.1160/TH08-12-0832. PubMed PMID:19652876.

34. Serres E, Debarbieux F, Stanchi F, Maggiorella L, Grall D, Turchi L, et al. Fibronectin expression in glioblastomas promotes cell cohesion, collective invasion of basement membrane in vitro and orthotopic tumor growth in mice. Oncogene. 2014;33(26):3451–62. doi:10.1038/onc.2013.305. PubMed PMID:23912459.

35. Ohnishi T, Hiraga S, Izumoto S, Matsumura H, Kanemura Y, Arita N, et al. Role of fibronectin-stimulated tumor cell migration in glioma invasion in vivo: clinical significance of fibronectin and fibronectin receptor expressed in human glioma tissues. Clin Exp Metastasis. 1998;16(8):729-41. PubMed PMID:10211986.

36. Neri D, Bicknell R. Tumour vascular targeting. Nat Rev Cancer. 2005;5(6):436–46. doi:10.1038/nrc1627. PubMed PMID:15928674.

37. Borsi L, Balza E, Bestagno M, Castellani P, Carnemolla B, Biro A, et al. Selective targeting of tumoral vasculature: comparison of different formats of an antibody (L19) to the ED-B domain of fibronectin. Int J Cancer. 2002;102(1):75–85. doi:10.1002/ijc.10662. PubMed PMID:12353237.

38. Peiris PM, Bauer L, Toy R, Tran E, Pansky J, Doolittle E, et al. Enhanced Delivery of Chemotherapy to Tumors Using a Multicomponent Nanochain with Radio-Frequency-Tunable Drug Release. ACS Nano. 2012;6(5):4157-68. Epub 2012/04/11. doi:10.1021/nn300652p. PubMed PMID:22486623.

39. Baumann BC, Kao GD, Mahmud A, Harada T, Swift J, Chapman C, et al. Enhancing the efficacy of drug-loaded nanocarriers against brain tumors by targeted radiation therapy. Oncotarget. 2013;4(1):64–79. doi:10.18632/oncotarget.777. PubMed PMID:23296073; PubMed Central PMCID:PMCPMC3702208.

